# Functional analysis of the N-terminal region of *Vibrio* FlhG, a MinD-type ATPase in flagellar number control

**DOI:** 10.1101/2022.03.19.484964

**Authors:** Michio Homma, Akira Mizuno, Yuxi Hao, Seiji Kojima

## Abstract

GTPase FlhF and ATPase FlhG are two key factors involved in regulating the flagellum number in *Vibrio alginolyticus*. FlhG is a paralog of the *Escherichia coli* cell division regulator MinD, which has a longer N-terminal region. Deleting the N-terminal region of FlhG results in cells with multiple flagella. The Q9A mutation in the DQAxxLR motif of the N-terminal region prevents it from activating the GTPase activity of FlhF *in vitro* and results in a multi-flagellation phenotype. The mutant FlhG protein was remarkably reduced compared to that of the wild-type protein *in vivo*. When the mutant FlhG was expressed at the same level as the wild-type FlhG, the number of flagella was restored to the wild-type level. Once synthesized in *Vibrio* cells, the N-terminal region mutation in FlhG seems not to affect the protein stability. We speculated that the *flhG* translation efficiency is decreased by N-terminal mutation. Our results suggest that the N-terminal region of FlhG controls the number of flagella by adjusting the FlhF activity and the amount of FlhG *in vivo*. We speculate that the regulation by FlhG, achieved through transcription by the master regulator FlaK, is affected by the mutations, resulting in reduced flagellar formation by FlhF.

## Introduction

Many motile bacteria can swim by rotating the flagellum which is a locomotive organ. The position and the number of flagella differ among bacterial species. To regulate the position and the number of flagella, many bacterial species use activation cycles of a MinD-type ATPase FlhG (also known as YlxH/FleN) and an SRP-FtsY-type GTPase FlhF (*1, 2*). The marine bacterium *Vibrio alginolyticus* has a flagellum at one cell pole, and the biogenesis of this single polar flagellum is regulated by FlhF and FlhG (*3*). The overexpression of FlhF in *V. alginolyticus* results in a phenotype of multiple polar flagella, whereas the depletion of FlhF results in a non-flagellated phenotype. Thus, FlhF is considered a positive factor in controlling the flagellar number. In contrast, overexpression of FlhG results in a non-flagellated phenotype, and its depletion results in a phenotype of multiple polar flagella. Thus, FlhG is considered a negative factor in controlling the flagellar number.

FlhF and FlhG are also known as transcriptional activators and repressors, respectively (*4, 5*). The polar flagellar genes of *Vibrio* form a regulon consisting of four hierarchies, from class 1 to class 4 operons (*6*). Expression of the class 2 gene group is controlled by FlaK (called FlrA in *V. cholerae* and FleQ in *Pseudomonas aeruginosa*), which functions as a master regulator and belongs to class 1 genes (Fig. S1). Both *flhF* and *flhG* are class 2 genes, and they are encoded by the same operon. However, FlhF acts as a transcription factor for class 3 genes, whereas FlhG inhibits class 2 gene transcription by FlaK in the presence of cyclic diguanylate (c-di-GMP). FlhG deficiency has been reported to increase flagellar gene expression (*4*).

In addition to flagellar gene regulation, FlhF, which is localized at the cell poles of *V. alginolyticus* (*7*), is involved in determining the location of flagellar formation. The *flhF* gene was first identified in *Pseudomonas putida* (*8*). There is a correlation between the polar localization of FlhF and flagellar formation. In *V. cholerae*, FlhF recruits FliF, a component of the MS-ring, to the inner membrane at the old cell pole in the early stage of flagellar formation (*9*). FlhF is a paralog of the signal recognition particle receptor FtsY, which functions together with Ffh, a signal recognition particle, in targeting nascent membrane proteins to the Sec translocon in the inner membranes of *Escherichia coli*. Both FtsY and Ffh, as well as FlhF, belong to the SRP-FtsY-type GTPase family and form an FtsY/Ffh heterodimer. FlhF also hydrolyzes GTP and forms a homodimer in the presence of GTP (*10*). After sending membrane protein precursors to the Sec transport machinery, the FtsY/Ffh heterodimer hydrolyzes GTP, becoming a GDP-bound form, and Ffh dissociates from FtsY. The GTPase activity of the FtsY/Ffh complex plays an important role in the processing of membrane proteins (*11, 12*). In *Bacillus subtilis*, the crystal structure of FlhF as a homodimer has been solved in the GTP-bound form (*11, 13*), although its function may not be the same as that of *Vibrio* FlhF because *B. subtilis* is not polar-flagellated. The GTPase activity of FlhF has also been shown in bacterial species, such as *Campylobacter* and *P. aeruginosa*, which have polar flagella, and binding to GTP and GTPase activity is important for FlhF function (*13–15*). The GTPase activity of FlhF increases in the presence of FlhG (*13, 16*). In our previous study on *V. alginolyticus* FlhF, polar flagellar formation and polar localization of FlhF were defective due to mutations in GTP binding and hydrolysis motifs (*7, 17*), indicating that GTPase motifs are important for the polar localization of FlhF. However, GTP involvement in this process remains unknown.

FlhG is a MinD paralog that constitutes the Min system together with MinC and MinE in *E. coli* and controls the Z ring formation composed of FtsZ at the cell division site. MinD is an ATPase, and its activity plays an important role in cell division. MinD forms a homodimer by binding ATP and binds to the membrane (*18, 19*). MinD binds MinC to promote depolymerization of FtsZ and activates MinC (*20*). The MinD/MinC complex dissociates from the membrane by MinE, and MinD and MinC move to another pole (*21*). During this process, MinE enhances the ATPase activity of MinD to hydrolyze the bound ATP (*22, 23*). By repeating such cycles in the Min system, it is thought that the position of the Z ring is determined, resulting in the formation of the cell division plate at the center of the cell. Cell division is defective when mutations are introduced into the ATPase binding site (K11, K16), the hydrolysis catalytic site (D40), and the activation site of ATPase (D152) (*24, 25*). FlhG has a conserved ATP motif, including the ATP-binding site K31 and K36 residues of MinD and D60 of the catalytic site for ATP hydrolysis. We have demonstrated that FlhG exhibits similar phenotypes in terms of ATPase activity in mutants corresponding to MinD mutations (*26*). However, the mutation of D171, corresponding to the ATPase activation site in MinD, resulted in a different phenotype: the ATPase activity of FlhG increased from 6- to 7-fold.

It has also been shown that FlhF and FlhG interact with each other (*7, 13*). Since the polar localization of FlhF decreases in the presence of FlhG, FlhG may play a role in retaining FlhF in the cytoplasm (*7*). The FlhG D171A mutant, which exhibits high ATPase activity, is strongly localized at the cell pole and severely inhibits flagellation. In contrast, FlhG mutants at putative ATP-binding sites (K31A, K36Q) abolished their polar localization and conferred a multi-flagellation phenotype. This suggests that FlhG at the cell pole is also involved in downregulating the flagellar number (*26*). The polar landmark membrane protein HubP controls the flagellar number in *V. alginolyticus* (*27*). Lack of HubP abolishes the polar localization of FlhG and causes the multi-flagellation phenotype, whereas polar localization of FlhF is not affected by *hubP* deficiency. FlhG is localized in a HubP-dependent manner and appears to directly inhibit the function of FlhF at the cell poles (*27*). Therefore, the positive control factor FlhF and the negative control factor FlhG are thought to determine the precise flagellar number by an exquisite balance of the activities with the assistance of appendage factors (*1*). It should be noted that *hubP* deficiency does not cause multi-flagellation in *Vibrio cholerae.* HubP function is, therefore, not always the same in *Vibrio* spp. (*1*).

The most striking difference between MinD and FlhG is in the most N-terminal region, which has ~ 20 additional residues of FlhG (Fig. 1A). N-terminal region contains a conserved motif, called DQAxxLR, in various bacterial species (Fig. 1B). FlhG stimulates FlhF-GTPase activity. However, mutations in the DQAxxLR motif of *B. subtilis* and *Campylobacter* FlhG abolish this effect (*13, 16*). Moreover, in *Campylobacter*, it has been shown that mutation in the DQAxxLR motif causes multiple flagellar phenotypes. Combined with the results of these previous studies and the finding that FlhF-GTPase activity is involved in the polar localization of FlhF and controlling the flagella-forming ability, the DQAxxLR motif appears to be involved in the regulation of the number of flagella. In this study, we investigated how the unique N-terminal sequence of *V. alginolyticus* FlhG is involved in regulating the flagellar number and multi-flagellation phenotype.

**Fig. 1.**
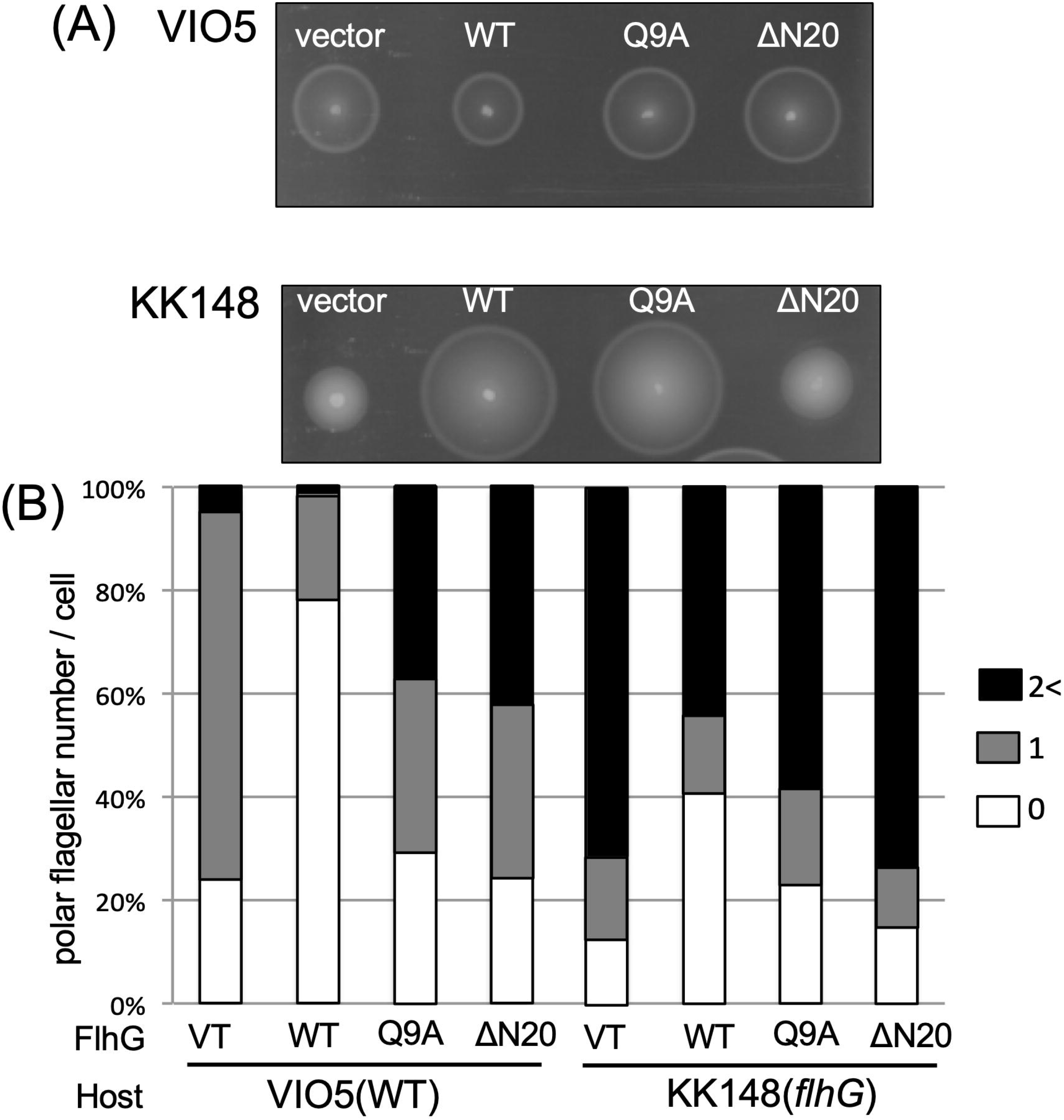
(A) FlhG is a homolog of MinD, and it has ATPase activity and conserved residues important for this activity. There is a unique N-terminal region in FlhG. Ec and Va represent *Escherichia coli* and *Vibrio alginolyticus*, respectively. The primary structure of FlhG is indicated. The N-terminal region of *V. alginolyticus* FlhG, which contains the DQAxxLR motif, lacks *E. coli* MinD. (B) Alignment of N-terminal FlhG and MinD. Many species have the DQAxxLR motif in the N-terminus of FlhG, and the glutamine residue at the position 9 of *V. alginolyticus* FlhG is highly conserved.

## Materials and Methods

### Strains, culture medium, and culture conditions

The strains and plasmids used in the present study are listed in Table S1. *E. coli* were grown in LB medium [1% (w/v) bactotryptone, 0.5% (w/v) yeast extract, and 0.5% (w/v) NaCl] at 37°C. For growth on a solid medium, 1.25% (w/v) agar was added to the LB medium. Wherever necessary, the antibiotics chloramphenicol and ampicillin were added to a final concentration of 25 μg/mL and 100 μg/mL, respectively, for *E. coli*. *V. alginolyticus* was cultured at 30°C in VC [0.5% (w/v) hipolypeptone, 0.5% (w/v) yeast extract, 0.4% (w/v) K_2_HPO_4_, 3% (w/v) NaCl, and 0.2% (w/v) glucose] or VPG medium [1% (w/v) hipolypeptone, 0.4% (w/v) K_2_HPO_4_, 3% (w/v) NaCl, and 0.5% (w/v) glycerol]. For growth on a solid medium, 1.25% (w/v) agar (Nakarai tesque) was added to the VC medium. To measure the motility in soft agar plates, 0.25% Bacto agar (Difco) was added to the VPG medium. Wherever necessary, chloramphenicol was added to a final concentration of 2.5 μg/mL for *V. alginolyticus*. The gene under the arabinose promoter (pBAD) control was induced by adding a suitable amount of arabinose for the intended expression level.

### DNA manipulations, mutagenesis, and sequencing

Routine DNA manipulations were conducted according to standard procedures. Mutations in *flhG* were introduced into plasmids by the “QuikChange” site-directed mutagenesis method, as described by Stratagene, and each mutation was confirmed by DNA sequencing. To construct the *flhG* deletion strain (NMB363) and the *flhF-egfp* chromosomal expression strain (NMB343), manipulation of chromosomal *flhF* and *flhG* was conducted using the pSW7848-based suicide vector and its conjugational transfer to the polar flagellar wild-type strain VIO5, as described previously (*28*). The nucleotide sequences for each *flhG* mutant were verified by DNA sequencing using a BigDye Terminator v3.1 Sequencing Standard Kit, performed by the Nagoya University DNA sequencing core facility. Transformation of *Vibrio* was carried out by electroporation using a Gene Pulser from Bio-Rad, as described previously (*29*).

### Western blotting

Each fraction separated by a size exclusion column was mixed with sodium dodecyl sulfate-polyacrylamide gel electrophoresis (SDS-PAGE) loading buffer and then heated to 95°C for 5 min. Proteins were separated by SDS-PAGE and then transferred onto polyvinylidene difluoride (PVDF) membranes. The membranes were incubated with anti-FlhG (FlhG B0728; *26*) and anti-flagellin (anti-PF45; *30*) primary antibodies. Horseradish peroxidase (HRP)-labeled rabbit anti-IgG antibody was used as the secondary antibody, diluted 1:10,000 in Tris-buffered saline (TBS)-Tween 20 (20 mM Tris-HCl [pH 7.5], 0.5 M NaCl, and 0.1% Tween 20). The enhanced chemiluminescence (ECL) reaction solution was then added, and the protein bands were detected using LAS 3000 (Fujifilm).

### Flagella detection by high-intensity dark-field microscopy

Overnight cultures were diluted 1:100 in VPG medium containing 0.02% or 0.2% (w/v) arabinose and 2.5◻μg/mL chloramphenicol and cultured at 30°C for a further 4 h. The cultured cells were harvested and suspended in V buffer (50 mM Tris-HCl (pH7.5), 300 mM NaCl, and 5 mM MgCl_2_). Flagella were observed under a high-intensity dark-field microscope (Olympus model BHT) equipped with a 100 W mercury lamp (Ushio USH-102).

### Fluorescence microscopy

*Vibrio* cells were cultured overnight in the VC medium at 30°C. The overnight cultures were diluted 1:100 in fresh VPG medium containing 0.02% (w/v) arabinose and 2.5◻μg/mL chloramphenicol and cultured for a further 4 h at 30°C. The cells were collected by centrifugation and suspended in V buffer at 1/4^th^ the volume of the original culture. Fluorescence microscopy was performed, as previously described (*26*).

### Purification of FlhG and FlhF

His-tagged FlhG was purified from BL21 (DE3)/pLysS cells containing the plasmid pTrc-*flhG*, grown in super broth (SB) medium containing ampicillin at 37°C with isopropyl ß-D-1-thiogalactopyranoside (IPTG) induction, as described previously (*26*). The concentrations of purified His-FlhG (mutant or wild-type) were determined by measuring the absorbance at 280 nm using an extinction coefficient of 11,460 (M^−1^ cm^−1^).

His-tagged FlhF was purified from *E. coli* BL21 (DE3) cells containing the plasmid pTSK110, grown at 37°C in LB medium containing ampicillin until OD_660_ of 0.5, then at 16°C in the presence of 0.4 mM IPTG overnight, as described previously (*10*). Aliquots of the eluted samples were frozen in liquid nitrogen and stored at −80°C. The concentration of FlhF was calculated from a calibration curve prepared by SDS-PAGE with a known concentration of BSA and quantification of the Coomassie brilliant blue (CBB)-stained bands using ImageJ software.

### Measurement of FlhG-ATPase activity

The ATPase activity of purified FlhG was measured using an ATPase assay kit (Innova Biosciences), as described previously (*26*). The purified FlhG was diluted with buffer A (20 mM Tris-HCl [pH 8.0], 300 mM NaCl, and 10% [w/v] glycerol) containing 65 mM imidazole. The diluted product, 50 μL, was mixed with 50 μL of buffer A containing 1 mM ATP and 10 mM MgCl_2_ in a 96-well plate and incubated at 25°C for 30 minutes. Next, 25 μL of a reaction terminator was added, and 2 min later, 10 μL of stabilizer was added, and the mixture was allowed to sit for a further 30 min at 25°C. The absorbance at 595 nm was measured using a plate reader (DS Pharma Biomedical). The concentration of inorganic phosphoric acid released into the solution was calculated from a calibration curve obtained using the absorbance measured for inorganic phosphoric acid of known concentration.

### Measurement of the FlhF-GTPase activity

GTPase activity was measured using a PiColorLock™ Gold Phosphate Detection System (Innova Bioscience), as described previously (*10*). Forty microliter samples of purified FlhF (8.5 μM) and FlhG (8.5 μM) were mixed with 40 μL of FlhF buffer [20 mM Tris-HCl (pH 8.0), 300 mM NaCl, 10 mM MgCl_2_, and 10 mM KCl] containing 1 mM GTP and then incubated for 30 min at 25°C. Next, 30 μL of the reaction terminator was added, and after 2 min, 12 μL of the stabilizer was added. After incubation at 25°C for 30 min, the absorbance at 595 nm was measured using a plate reader (DS Pharma Biomedical). The concentration of inorganic phosphoric acid was calculated from a calibration curve obtained using inorganic phosphate at known concentrations.

## Results

### Roles of the N-terminal region of FlhG for flagellar number regulation

To clarify the function of the N-terminal region of FlhG, an N-terminal 20-residue deletion mutant of FlhG (ΔN20) and an amino acid substitution mutant (Q9A) were constructed (Fig. 1). *V. alginolyticus* exhibits a multi-flagellation phenotype at the cell poles in response to *flhG* deficiency, and a non-flagellate phenotype in response to overexpression of FlhG (*3*). Motility was evaluated by the size of the motility rings formed on soft agar plates containing 0.02% arabinose (Fig. 2A). *V. alginolyticus* spotted in the soft agar medium and grown for several hours was observed to swim in the medium. If the cells do not have flagella, the ring does not expand. When the cells have multiple flagella, the flagella are entangled with each other. As this restricts swimming, the ring size is smaller than when the cells have a single flagellum. Expression of wild-type FlhG from the arabinose-inducible plasmid in VIO5, which is the wild-type strain for polar flagellar formation, decreased the swimming ability compared to those of the vector control strain. When the ΔN20-mutant and Q9A-mutant FlhG were expressed by their respective plasmids, the swimming abilities were similar to those of the vector control strain. Expression of wild-type FlhG by the plasmid improved motility of the *flhG* mutant strain KK148. However, expression of ΔN20 FlhG had almost no effect on motility and exhibited a ring size similar to that of the original mutant cells (vector). Expression of the Q9A-mutant FlhG in KK148 conferred motility similar to that of the wild-type (WT) strain. Thus, the N-terminal region specific to FlhG plays an important role in reducing the flagellar number.

**Fig. 2.**
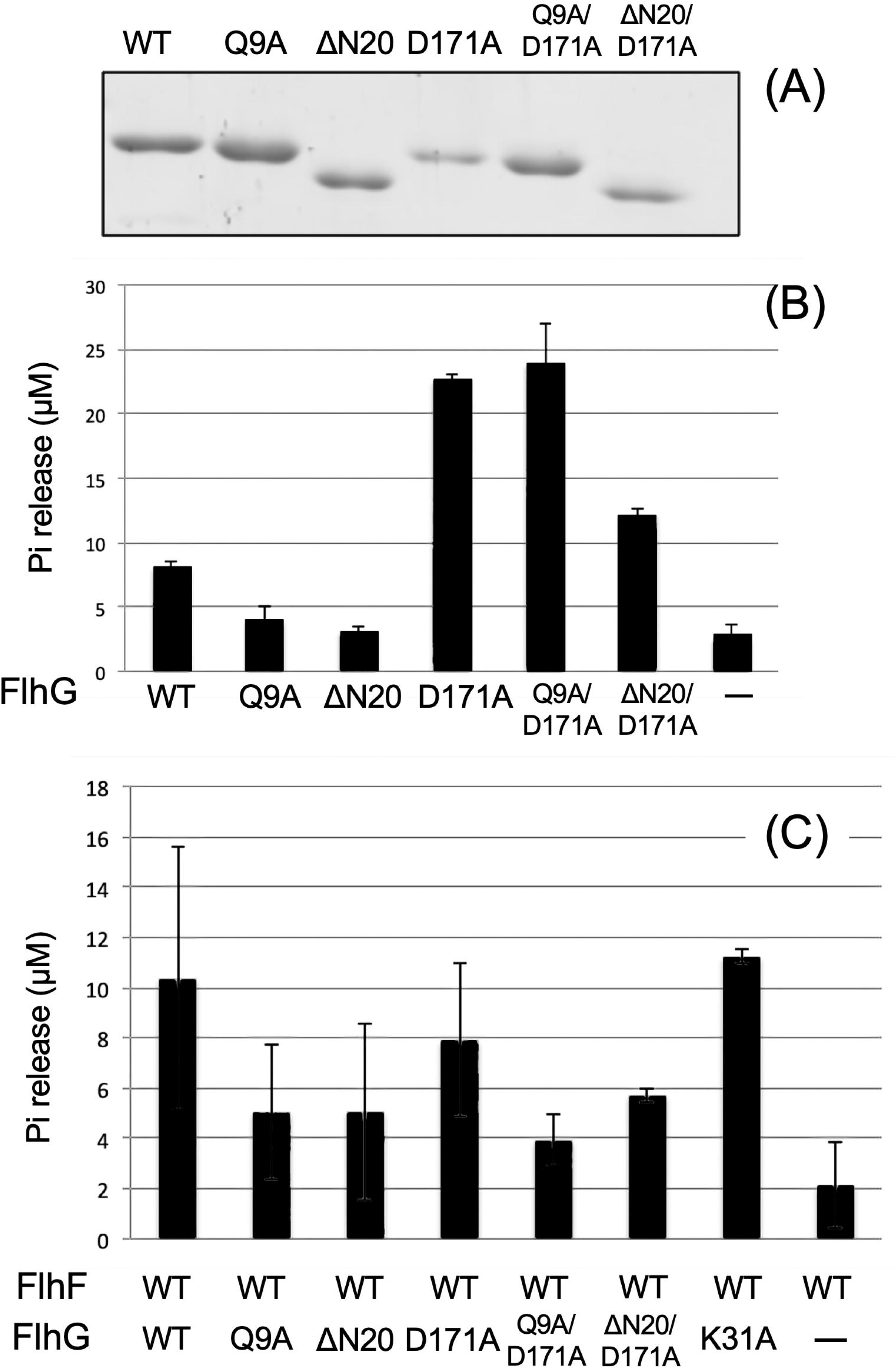
Motility and flagellar number of bacterial cells. VIO5 (wild-type strain) and KK148 (the *flhG*-deficient strain) containing pBAD33 as vector control and pAK520 encoding *flhG* were cultured overnight at 30°C in VC medium containing 0.02% arabinose. (A) A 2 μL aliquot of the overnight culture was spotted on a VPG agar plate containing 0.02% arabinose and incubated at 30°C for 7 h (B). The overnight culture was diluted 100-fold in VPG medium containing 0.02% and cultured at 30°C for 4 h. Independent experiments were conducted three times to observe more than 100 cells using a high-intensity dark-field microscope.

Next, we examined the flagellar number of the cells (Fig. 2B). When the empty plasmid (vector) was introduced into the wild-type VIO5 strain and cultured in the presence of 0.02% arabinose, 72% of the cells formed a single flagellum at the cell poles, and 24% of the cells had no flagella. When a plasmid containing wild-type FlhG was introduced into the VIO5 strain and induced with 0.02% arabinose, 22% of the cells formed a single flagellum, and 77% of the cells had no flagella. In contrast, when a plasmid containing the ΔN20 or the Q9A FlhG mutant was introduced and then induced with 0.02% arabinose, approximately 34% and 32%, respectively, of the cells formed a single flagellum and approximately 23% and 32%, respectively, of the cells had no flagella. In addition, we found that the number of cells with multiple flagella increased only when ΔN20 or Q9A were produced in the cells. The same experiments were performed using the *flhG-deficient* strain KK148. When FlhG was not expressed by a vector, approximately 72% of the cells had multiple flagella, and approximately 13% had no flagella. When wild-type FlhG was expressed, approximately 44% of the cells had multiple flagella, and approximately 40% had no flagella. When ΔN20 or Q9A FlhG were expressed, approximately 74% and 59%, respectively, of the cells had multiple flagella, while approximately 15% and approximately 20%, respectively, of the cells had no flagella. Based on these results, the ΔN20 FlhG mutant likely suppresses wild-type FlhG function.

### Mutations of the N-terminal region of FlhG involved in FlhF-GTPase activity

In various bacterial species, the DQAxxLR sequence motif is widely conserved in the N-terminal region of FlhG (Fig. 1B). Previous studies on *Bacillus subtilis* and *Campylobacter* have shown that FlhF-GTPase activity increases in the presence of FlhG, but not by FlhG with the Q4A mutation (*13, 16*). Therefore, we purified recombinant FlhF from *V. alginolyticus* and assayed its GTPase activity in the presence of wild-type or mutant FlhG protein (Fig. 3). For purification, a His-tag was attached to FlhG at the N-terminus or to FlhF at the C-terminus. FlhF and FlhG were adjusted to the same concentration in the reactions, and GTP hydrolysis was then measured (Fig. 3C). In the presence of wild-type FlhG, ~10 μM of inorganic phosphate was released by FlhF. When the GTPase activity of FlhF alone was measured by adding a buffer, approximately 2 μM of inorganic phosphate was released. Thus, the FlhF-GTPase activity increased in the presence of FlhG, as in other bacteria. In the presence of the FlhG Q9A or ΔN20 mutant, approximately 5 μM inorganic phosphate was released. This suggests that the N-terminal region of FlhG enhances FlhF-GTPase activity.

**Fig. 3.**
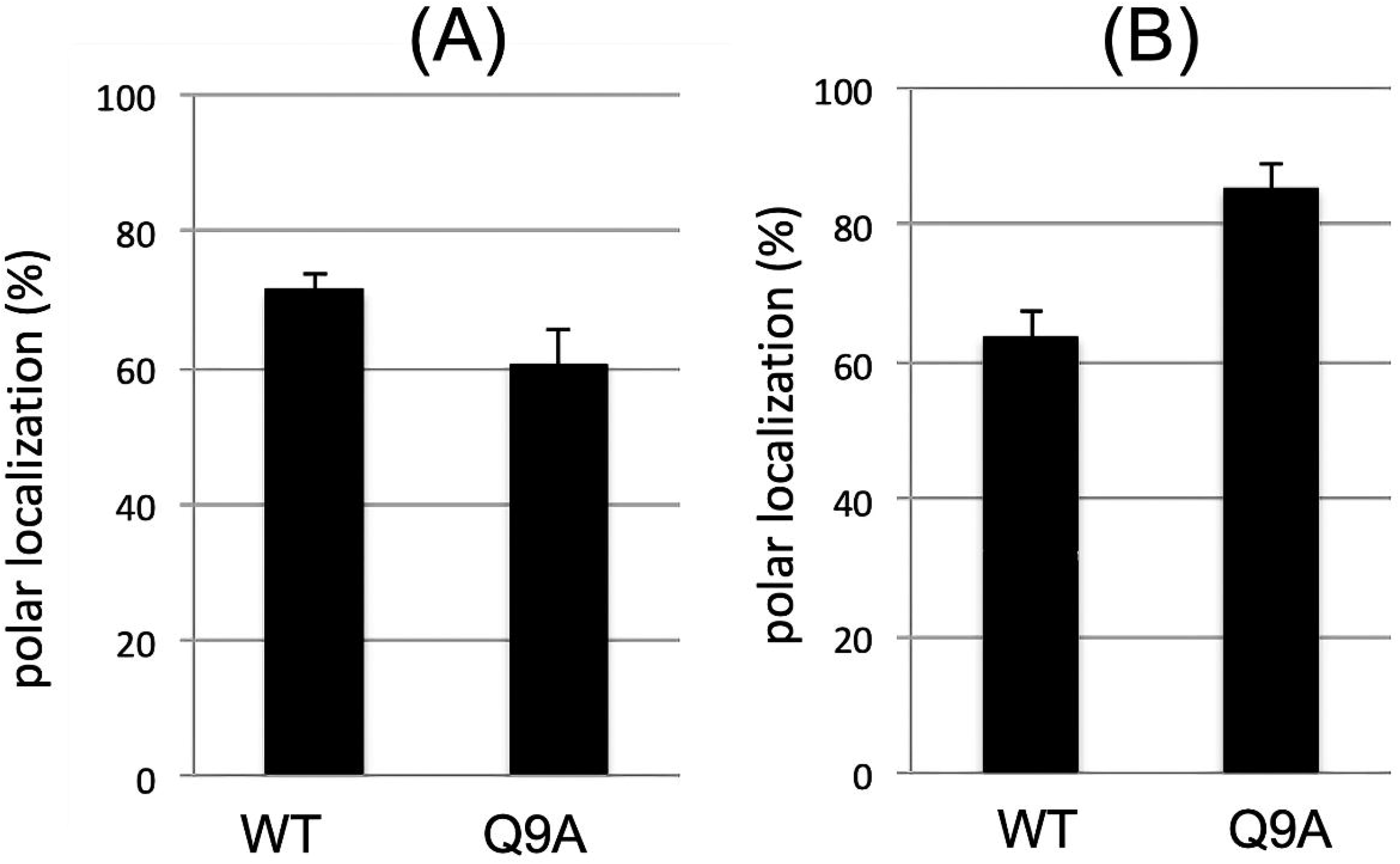
ATPase activity of FlhG. (A) The FlhG proteins were prepared from BL21(DE3)/pLysS cells harboring a plasmid encoding pTrc-flhG (WT) or mutant derivatives and purified by His-tag affinity. The concentration of each purified FlhG was calculated from the absorbance at 280 nm and adjusted to 10 μM with buffer A containing 62.5 mM imidazole. The proteins (5 μL aliquots) were separated by SDS-PAGE and stained with CBB. (B) FlhG-ATPase activity measurement. The release of inorganic phosphoric acid was measured after mixing 10 μM purified FlhG with 1 mM ATP and 10 mM MgCl_2_ and incubation for 30 min. (C) FlhF-GTPase activity measurement. The release of inorganic phosphoric acid was measured after mixing 8.5 μM purified FlhF with 8.5 μM purified FlhG, 1 mM GTP, and 10 mM MgCl_2_ and incubation for 30 min.

### Properties of FlhG-ATPase activity mutants

Wild-type FlhG on its own is known to have low ATPase activity. Sequence alignment predicts that K31 and K36 of *V. alginolyticus* FlhG are in the ATP binding site, D60 is a catalytic residue, and D171 is essential for activation of ATPase activity. K31A and D60A mutations abolish FlhG-ATPase activity, while the D171A mutation increases the ATPase activity six to seven times (*26*). We investigated whether the Q9A/D171A and ΔN20/D171A double mutations in the N-terminal region of FlhG affect FlhG-ATPase activity. The concentration of purified FlhG was adjusted to 10 μM, and the ATP hydrolysis activity was then measured. Wild-type FlhG released approximately 8 μM inorganic phosphate, whereas FlhG Q9A and ΔN20 mutants released approximately 4 μM and 3 μM inorganic phosphate, respectively (Fig. 3B). This suggests that N-terminal mutations reduce the FlhG-ATPase activity. In contrast, when combined with the D171A mutation, the Q9A/D171A mutant released approximately 23 μM inorganic phosphate, comparable to that released by the D171A mutant. The FlhG-ATPase activity was not affected by the Q9A mutation in the D171A mutant. The ΔN20/D171A double mutant released approximately 12 μM inorganic phosphate, approximately half of that released by the D171A mutant. Thus, the Q9A and ΔN20 mutations affect the ATPase activity of FlhG but do not or only slightly affect the high activity of the D171A mutant. The N-terminal region may not directly participate in the ATPase activity of FlhG.

We examined the effects of mutant D171A with high ATPase activity and the Q9A/D171A and ΔN20/D171A double mutants of FlhG on FlhF-GTPase activity (Fig. 3C). In the presence of the D171A mutant, it released approximately 8 μM of inorganic phosphate, indicating an increase in GTPase activity, albeit not as high as that of the wild type. The FlhF-GTPase activity did not increase in the presence of either of the double mutants. Mutation of K31A, which is in the ATP binding site of FlhG, resulted in a loss of ATPase activity (*26*). The FlhF-GTPase activity increased in the presence of K31A FlhG, similar to that of wild-type FlhG.

### The polar localization ability of FlhG

The polar localization ability of FlhG-GFP was examined in the *flhG* mutant of KK148 containing the pAK541 plasmid coding for *flhG-egfp*. Polar-localized dots were observed in 72% of the cells expressing FlhG-GFP, whereas Q9A FlhG had a polar localization in approximately 60% of the cells (Fig. 4). The polar localization of FlhF was examined using NMB343, which was constructed to express chromosomal *flhF-egfp*. When wild-type FlhG was expressed by the plasmid in NMB343, FlhF-GFP had a polar localization in approximately 64% of cells, which increased to approximately 86% by Q9A mutation of FlhG.

**Fig. 4.**
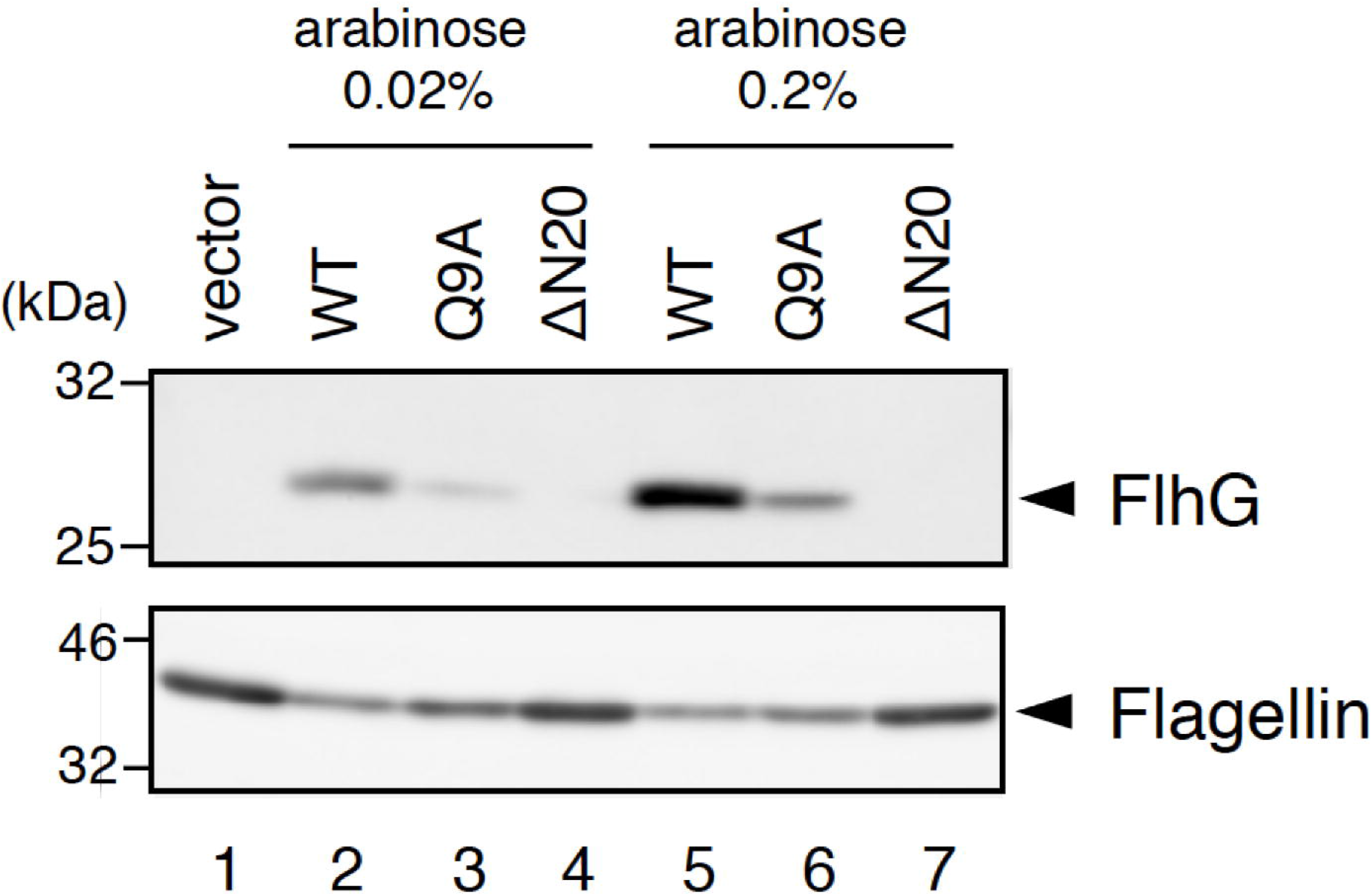
The polar localization of FlhF and FlhG in *Vibrio* cells. (A) KK148 containing pAK541 encoding wild-type or Q9A *flhG-egfp* was cultured at 30°C for 4 h in a VPG medium containing 0.02% arabinose. (B) NMB343, genetically engineered to express chromosomal *flhF-egfp*, containing pAK520 encoding *flhG* (WT) or Q9A mutant *flhG* (Q9A), was cultured in a VPG medium containing 0.02% arabinose at 30°C for 4 h. These samples were subjected to three independent experiments using a fluorescence microscope to count the number of cells in which fluorescent dots were observed at the poles.

Next, the amount of FlhG in the *flhG* mutant strain KK148 was examined by western blotting (Fig. 5). The expression level of the Q9A mutant was lower than that of the wild-type protein, and ΔN20 FlhG was not detected in the cells. The amount of flagellin in the cells was increased by the expression of Q9A FlhG and ΔN20 FlhG compared to that of the wild-type FlhG. Thus, the loss of FlhG function *in vivo* caused by N-terminal mutation stems from a decrease in FlhG production.

**Fig. 5.**
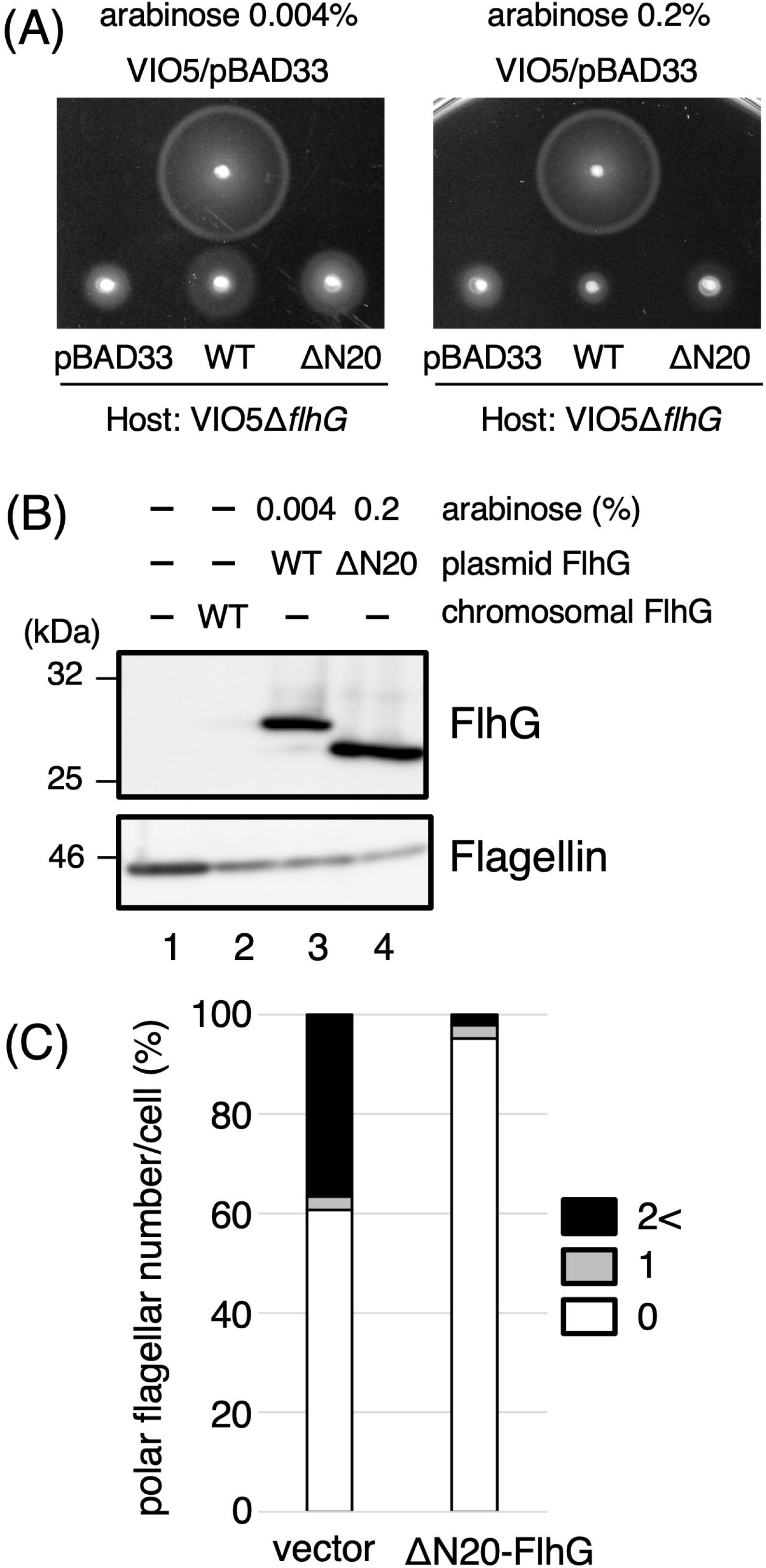
Protein expression levels of wild-type and mutant FlhG. KK148 containing pAK520 encoding wild-type *flhG* (WT), Q9A mutant (Q9A), or ΔN20 mutant (ΔN20), was cultured at 30°C for 4 h in VPG medium containing 0.02% or 0.2% arabinose. The whole-cell proteins were separated by SDS-PAGE, and the blotted membranes were probed with anti-FlhG or anti-Flagellin antibodies.

### The effect of wild-type and mutant FlhG expression in vivo

To verify the above possibility, we increased the expression level of ΔN20 FlhG in *Vibrio* cells using the overexpression plasmid pTSK151, in which *flhG* is joined to an optimized Shine-Dalgarno sequence (Fig. 6). Using this construct, *flhG* expression could be adjusted by alteration of the arabinose concentration in the medium. Since KK148 carries a nonsense mutation in the middle of *flhG* (Q109amber), and to eliminate complexity by the heterogeneity of the FlhG mutant protein, we deleted *flhG* from the VIO5 strain. This *flhG*-deletion strain (NMB363) exhibited a multipolar flagellar phenotype similar to that of strain KK148. NMB363 exhibited reduced motility compared to wild-type VIO5 on soft agar plates (Fig. 6A, vector control pBAD33). The motility was only slightly restored when wild-type and ΔN20 FlhG were expressed by the plasmid pTSK151 and induced by 0.004% arabinose (left panel). In contrast, when wild-type and ΔN20 FlhG were overproduced by 0.2% arabinose (right panel), their motility was even lower than that of the vector control, indicating that flagellation was strongly inhibited, even by ΔN20 FlhG.

**Fig. 6.**
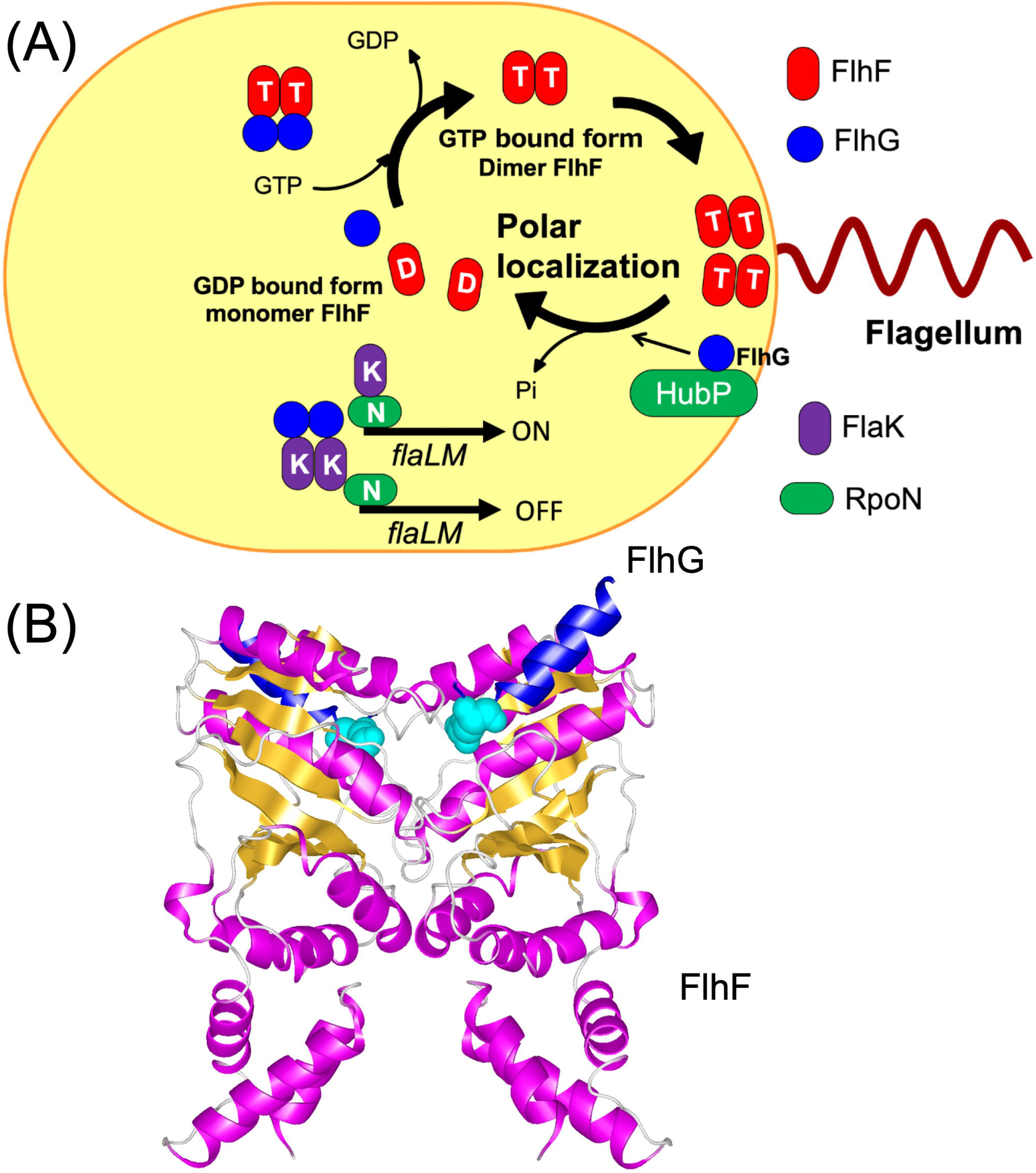
The effect of the expression level of wild-type and mutant FlhG *in vivo*. (A) Overproduction of ΔN20 FlhG mutant inhibited the motility of cells in soft agar plates. The strain VIO5 (upper part of the plate) or NMB363(VIO5Δ*flhG*) (lower part of the plate) was transformed with the empty vector pBAD33, or pTSK151 expressing wild-type FlhG (WT) or ΔN20 FlhG (ΔN20). Overnight cultures of each strain in the VC medium were spotted on the VPG soft agar plate containing 0.004% (left panel) or 0.2% (right panel) arabinose and incubated at 30°C for 5 h. (B) Immunoblot of the whole-cell lysates of overproduced wild-type FlhG or ΔN20 FlhG. Each strain was cultured in a VPG medium with the indicated concentrations of arabinose for 3 h at 30°C, and immunoblotting was performed with anti-FlhG (upper panel) or anti-Flagellin (lower panel) antibodies. Lane 1, NMB363/pBAD33 no arabinose; Lane 2, VIO5/pBAD33 no arabinose; Lane 3, NMB363/pTSK151(WT) induced with 0.004% arabinose; Lane 4, NMB363/pTSK151(ΔN20) induced with 0.2% arabinose. (C) The polar flagellar number of NMB363 cells overproducing ΔN20 FlhG. NMB363 cells harboring the empty plasmid pBAD33 (vector) or pTSK151 encoding Δ*N20-flhG* mutant (ΔN20 FlhG) were cultured in VPG medium at 30°C for 4 h. ΔN20 FlhG was induced with 0.2% arabinose. Cells were observed by high-intensity dark-field microscopy, and the polar flagella of each cell were counted for at least 700 individual cells. The results shown here were obtained from duplicate experiments. The percentage of the cells with more than two polar flagella is shown by filled bars, cells with one polar flagellum are shown by gray bars, and cells with no flagellum are shown by open bars.

Protein production was confirmed by western blotting analysis of whole-cell lysates of these strains (Fig. 6B). In the presence of 0.2% arabinose, the level of ΔN20 FlhG protein (lane 4) was comparable to the level of wild-type FlhG produced by 0.004% arabinose (lane 3). Overexpression of ΔN20 FlhG also reduced flagellin levels (lane 4, lower panel). Consistent with these results, flagellation was severely inhibited by overexpression of ΔN20 FlhG by pTSK151. Approximately 36% of the NMB363 cells with vector plasmid pBAD33 had two or more flagella, and 60% had no flagella. On the other hand, only approximately 2% of the NMB363 cells overproducing ΔN20 FlhG had two or more flagella, and 95% of them had no flagella (Fig. 6C). These results suggest that the N-terminal region of FlhG is involved in FlhF-GTPase activity. However, the mutation caused a multi-flagellated phenotype mainly due to the decrease in the amount of FlhG *in vivo*. Next, we examined whether the N-terminal mutations affected the stability of FlhG. After the induction of FlhG expression by arabinose, cell growth and protein production were inhibited by the addition of kanamycin, and the cells were sampled to investigate the cellular level of FlhG. We confirmed that in both the wild-type and Q9A FlhG strains, the protein levels did not change, suggesting that the proteins are stable under *in vivo* conditions (Fig. S2).

## Discussion

### Role of the DQAxxLR motif in V. alginolyticus

FlhF, which positively regulates the flagellum number, is a paralog of the SRP-type GTPase FtsY/Ffh, and all GTPase motifs are conserved. The polar localization of FlhF is necessary for flagellar formation, and the GTPase motif is important for the polar localization (*1*). Therefore, FlhF-GTPase activity is thought to be involved in controlling the flagellum number. As mentioned above, FlhG is a paralog of the MinD ATPase, which is a cell division surface regulator. The characteristic N-terminal region of FlhG contains the conserved DQAxxLR motif. In *B. subtilis* and *Campylobacter*, FlhF-GTPase activity is enhanced in the presence of FlhG, and this effect is lost by mutation of the DQAxxLR motif (*13, 16*). Furthermore, in *Campylobacter*, the Q4A mutation in this motif conferred the same phenotype as *flhG* deletion, resulting in many flagella at the cell pole. Our results show that *Vibrio* FlhG has biochemical properties similar to those of other species (Fig. 3). Q9A FlhG and ΔN20 FlhG did not increase the FlhF-GTPase activity, suggesting that the N-terminal region of FlhG in *V. alginolyticus*, containing the DQAxxLR motif, is involved in the control of the FlhF-GTPase activity. The ATPase activity of FlhG was reduced but not abolished by these N-terminal mutations, suggesting that the N-terminal region is important for the activation of FlhG-ATPase activity.

Expression of Q9A FlhG and ΔN20 FlhG did not complement the *flhG*-deficient strain KK148 and conferred a multi-flagellation phenotype. We initially hypothesized that defects in the DQAxxLR motif impaired the negative flagellation activity of FlhG, but this does not appear to be the case. We found that the cellular levels of Q9A and ΔN20 FlhG were markedly reduced (Fig. 5). In particular, ΔN20 FlhG was below the detection level by western blotting analysis. Adjusting the protein levels of these mutants by increasing the arabinose concentration (for Q9A) or by optimizing the construct caused severe inhibition of flagellation (Fig. 6). Furthermore, the Q9A mutant can localize to the cell pole, similar to wild-type FlhG (Fig. 4). These results indicate that the DQAxxLR motif is dispensable for FlhG function in negative flagellar formation. Mutations in the DQAxxLR motif affect the ATPase activity of FlhG and the FlhG protein level.

### Control of the level of FlhG protein by the N-terminal region of FlhG

Protein levels are mainly controlled by two factors: synthesis and degradation. We found that the Q9A mutant proteins were as stable as wild-type FlhG (Fig. S2). Moreover, as the N-terminal His-tag stabilized the Q9A and ΔN20 FlhG proteins overexpressed in *E. coli*, we purified these mutants for subsequent biochemical analyses. Once the proteins are synthesized, they are refractory to degradation, and the N-terminal region did not appear to affect the stability of FlhG. We believe that the next possibility is that the N-terminal region regulates the amount of FlhG *in vivo* at the translational or transcriptional level. In transcription regulation, it has been thought that the *flhA*, *flhF*, *flhG*, and *fliA* genes form an operon, and according to studies of *V. cholerae* and *V. parahaemolyticus*, they are class 2 genes (Fig. S1) (*6*). The *flhA* operon is transcribed by FlaK, which is a master regulator of flagellum-related genes. FlhF promotes the transcription of class 3 genes, and FlhG suppresses the transcription of class 2 genes by FlaK (*5*). Reduction of the level of the N-terminal mutant FlhG proteins impaired the regulation of the transcription of class 2 genes by FlaK, which regulates the expression of other flagellum-related genes, including *flhF*. Thus, the cells may produce many flagella because the level of FlhF is increased. The *flhF* and *flhG* genes form an operon in the chromosome and are expected to be translated from the same mRNA. Based on the present results, we speculated that the translation efficiency of *flhG* is decreased by the N-terminal mutations.

### Model of the control of the flagellar number by V. alginolyticus FlhG

The model for flagellar number regulation based on the present and our previous studies (*1*) is shown in Fig. 7. The N-terminal region of FlhG appears to be involved in determining the number of flagella by affecting mRNA translation efficiency. The *flhG* gene transcribed by the plasmid promoter can be induced by arabinose. Thus, the mRNA levels should be the same for the wild-type and the mutant *flhG* genes. A large amount of FlhG expressed by the plasmid undergoes negative feedback control and inhibits transcriptional promotion by FlaK, which interacts with FlhG. Consequently, the mRNA levels of *flhA*, *flhF*, *flhG*, and *fliA* are reduced, and the cells, therefore, cannot generate flagella. If the translation efficiency of the mutant *flhG* mRNA decreases, there should be no negative feedback control. However, our biochemical analysis of the mutant proteins showed that the mutations reduce the ATPase activity of FlhG and the activation ability of FlhF GTPase. This model does not completely explain the results of our *in vivo* and *in vitro* data. To verify this model for the function of the N-terminal region, it is necessary to clarify whether there is a transcriptional activity of class 2 genes by FlaK, and interaction between FlhG and FlaK is yet to be demonstrated. Thus, the formation of a single polar flagellum is subject to complex regulatory mechanisms.

**Fig. 7.**
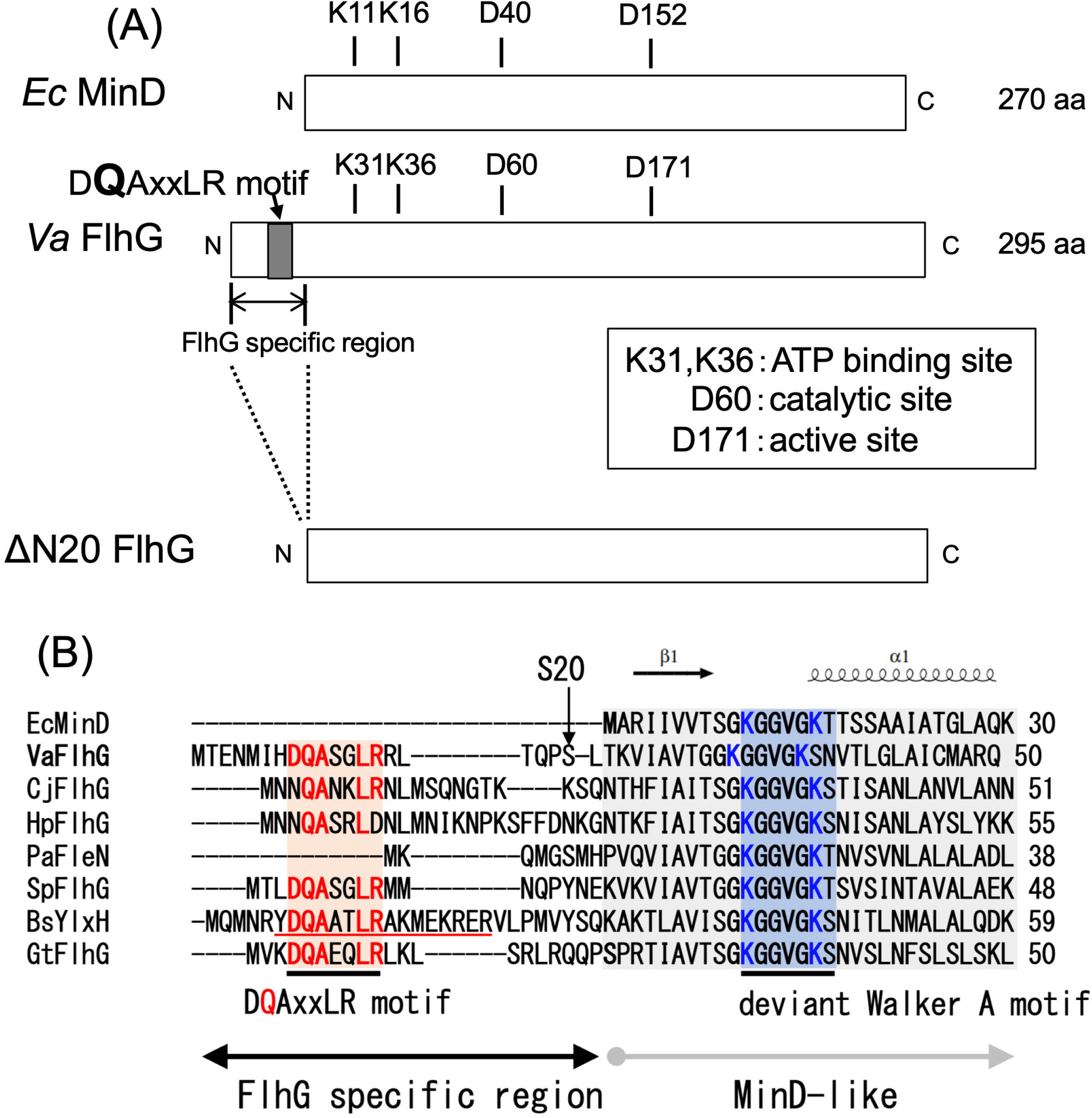
(A) The model of FlhF polar localization in *V. alginolyticus*. FlhF diffuses into the cytoplasm as a GDP-bound monomer, but when it binds to GTP, it forms a dimer and becomes localized at the pole. Localized FlhF promotes flagellar formation at the cell pole. FlhG diffuses in the cytoplasm and interacts with FlhF to inhibit polar localization of FlhF. At the poles, the GTPase activity of FlhF is activated by FlhG, which is localized at the pole by the HubP scaffold. FlhF is converted to the GDP-bound form and dissociates from the cell pole. In wild-type *V. alginolyticus*, the amount of FlhF localized at the pole is controlled to a level resulting in a single, polar flagellum biosynthesis. The figure is quoted from our previous paper (*10*). (B) The complex structure model of FlhF and N-terminal FlhG (YlxH). The crystal structure model of PDB 3SYN of *Bacillus subtilis* is shown in the helix model. The N-terminal region of FlhG (YlxH) is shown in blue, and the corresponding residue Q9 in *V. alginolyticus* is shown in the space-filling model in sky blue.

## Acknowledgments

This work was supported in part by the Japan Society for the Promotion of Science (JSPS) KAKENHI [Grant Number 20H03220 (to M.H.)].

## Supporting information

The supplementary information associated with this article can be found online on the publisher’s website.

## Figure legends

**Fig. S1.** Diagram showing the class of transcriptional control of flagellum-related factors in *V. alginolyticus*. FlhF and FlhG belong to class 2, FlhF promotes expression of class 3 gene, and FlhG inhibits the expression of class 2 genes by FlaK, which is a master regulator of flagellum-related factors.

**Fig. S2.** Protein stability of the FlhG mutants. KK148 containing pAK520 encoding wild-type *flhG* (WT) or Q9A mutant (Q9A) was cultured at 30 C in a VPG medium containing 0.02% arabinose. After 4 h, 1 mg/mL of kanamycin was added to the culture medium, and samples of the cells were collected every hour. The proteins in the cells were separated by SDS-PAGE, and the blotted membranes were probed with an anti-FlhG antibody.

## Notes

### Competing Interest Statement

The authors have declared no competing interest.

## References

(1) Kojima, S., Terashima, H., and Homma, M. (2020) Regulation of the single polar flagellar biogenesis. Biomolecules 10, 533

(2) Schuhmacher, J.S., Thormann, K.M., and Bange, G. (2015) How bacteria maintain location and number of flagella? FEMS Microbiol. Rev. 39, 812–822

(3) Kusumoto, A., Kamisaka, K., Yakushi, T., Terashima, H., Shinohara, A., and Homma, M. (2006) Regulation of polar flagellar number by the *flhF* and *flhG* genes in *Vibrio alginolyticus*. J. Biochem. 139, 113–121

(4) Correa, N.E., Peng, F., and Klose, K.E. (2005) Roles of the regulatory proteins FlhF and FlhG in the *Vibrio cholerae* flagellar transcription hierarchy. J. Bacteriol. 187, 6324–6332

(5) Kojima, M., Nishioka, N., Kusumoto, A., Yagasaki, J., Fukuda, T., and Homma, M. (2011) Conversion of mono-polar to peritrichous flagellation in Vibrio alginolyticus. Microbiol. Immunol. 55, 76–83

(6) Homma, M., Nishikino, T., and Kojima, M. (2022) Achievements in bacterial flagellar research with focus on *Vibrio* species. Microbiol. Immunol. 66, 75–95

(7) Kusumoto, A., Shinohara, A., Terashima, H., Kojima, S., Yakushi, T., and Homma, M. (2008) Collaboration of FlhF and FlhG to regulate polar-flagella number and localization in *Vibrio alginolyticus*. Microbiology 154, 1390–1399

(8) Pandza, S., Baetens, M., Park, C.H., Au, T., Keyhan, M., and Matin, A. (2000) The G-protein FlhF has a role in polar flagellar placement and general stress response induction in *Pseudomonas putida*. Mol. Microbiol. 36, 414–423

(9) Green, J.C., Kahramanoglou, C., Rahman, A., Pender, A.M., Charbonnel, N., and Fraser, G.M. (2009) Recruitment of the earliest component of the bacterial flagellum to the old cell division pole by a membrane-associated signal recognition particle family GTP-binding protein. J. Mol. Biol. 391, 679–690

(10) Kondo, S., Imura, Y., Mizuno, A., Homma, M., and Kojima, S. (2018) Biochemical analysis of GTPase FlhF which controls the number and position of flagellar formation in marine *Vibrio*. Sci. Rep. 8, 12115

(11) Bange, G., Petzold, G., Wild, K., Parlitz, R.O., and Sinning, I. (2007) The crystal structure of the third signal-recognition particle GTPase FlhF reveals a homodimer with bound GTP. Proc. Natl. Acad. Sci. USA 104, 13621–13625

(12) Cross, B.C., Sinning, I., Luirink, J., and High, S. (2009) Delivering proteins for export from the cytosol. Nat. Rev. Mol. Cell Biol. 10, 255–264

(13) Bange, G., Kummerer, N., Grudnik, P., Lindner, R., Petzold, G., Kressler, D., Hurt, E., Wild, K., and Sinning, I. (2011) Structural basis for the molecular evolution of SRP-GTPase activation by protein. Nat. Struct. Mol. Biol. 18, 1376–1380

(14) Balaban, M., Joslin, S.N., and Hendrixson, D.R. (2009) FlhF and its GTPase activity are required for distinct processes in flagellar gene regulation and biosynthesis in *Campylobacter jejuni*. J. Bacteriol. 191, 6602–6611

(15) Schniederberend, M., Abdurachim, K., Murray, T.S., and Kazmierczak, B.I. (2013) The GTPase activity of FlhF is dispensable for flagellar localization, but not motility, in *Pseudomonas aeruginosa*. J. Bacteriol. 195, 1051–1060

(16) Gulbronson, C.J., Ribardo, D.A., Balaban, M., Knauer, C., Bange, G., and Hendrixson, D.R. (2016) FlhG employs diverse intrinsic domains and influences FlhF GTPase activity to numerically regulate polar flagellar biogenesis in *Campylobacter jejuni*. Mol. Microbiol. 99, 291–306

(17) Kondo, S., Homma, M., and Kojima, S. (2017) Analysis of the GTPase motif of FlhF in the control of the number and location of polar flagella in *Vibrio alginolyticus*. Biophy. Physicobiol. 14, 173–181

(18) Hu, Z., Gogol, E.P., and Lutkenhaus, J. (2002) Dynamic assembly of MinD on phospholipid vesicles regulated by ATP and MinE. Proc. Natl. Acad. Sci. USA 99, 6761–6766

(19) Hu, Z., and Lutkenhaus, J. (2003) A conserved sequence at the C-terminus of MinD is required for binding to the membrane and targeting MinC to the septum. Mol. Microbiol. 47, 345–355

(20) Zhou, H., and Lutkenhaus, J. (2004) The switch I and II regions of MinD are required for binding and activating MinC. J. Bacteriol. 186, 1546–1555

(21) Hu, Z., and Lutkenhaus, J. (2001) Topological regulation of cell division in *E. coli*. spatiotemporal oscillation of MinD requires stimulation of its ATPase by MinE and phospholipid. Mol. Cell 7, 1337–1343

(22) Park, K.T., Wu, W., Battaile, K.P., Lovell, S., Holyoak, T., and Lutkenhaus, J. (2011) The Min oscillator uses MinD-dependent conformational changes in MinE to spatially regulate cytokinesis. Cell 146, 396–407

(23) Park, K.T., Wu, W., Lovell, S., and Lutkenhaus, J. (2012) Mechanism of the asymmetric activation of the MinD ATPase by MinE. Mol. Microbiol. 85, 271–281

(24) Ma, L., King, G.F., and Rothfield, L. (2004) Positioning of the MinE binding site on the MinD surface suggests a plausible mechanism for activation of the *Escherichia coli* MinD ATPase during division site selection. Mol. Microbiol. 54, 99–108

(25) Zhou, H., Schulze, R., Cox, S., Saez, C., Hu, Z., and Lutkenhaus, J. (2005) Analysis of MinD mutations reveals residues required for MinE stimulation of the MinD ATPase and residues required for MinC interaction. J. Bacteriol. 187, 629–638

(26) Ono, H., Takashima, A., Hirata, H., Homma, M., and Kojima, S. (2015) The MinD homolog FlhG regulates the synthesis of the single polar flagellum of *Vibrio alginolyticus*. Mol. Microbiol. 98, 130–141

(27) Takekawa, N., Kwon, S., Nishioka, N., Kojima, S., and Homma, M. (2016) HubP, a Polar Landmark Protein, Regulates Flagellar Number by Assisting in the Proper Polar Localization of FlhG in *Vibrio alginolyticus*. J. Bacteriol. 198, 3091–3098

(28) Kojima, S., Yoneda, T., Morimoto, W., and Homma, M. (2019) Effect of PlzD, a YcgR homologue of c-di-GMP-binding protein, on polar flagellar motility in Vibrio alginolyticus. J. Biochem. 166, 77–88

(29) Kawagishi, I., Okunishi, I., Homma, M., and Imae, Y. (1994) Removal of the periplasmic DNase before electroporation enhances efficiency of transformation in a marine bacterium *Vibrio alginolyticus*. Microbiology 140, 2355–2361

(30) Nishioka, N., Furuno, M., Kawagishi, I., and Homma, M. (1998) Flagellin-containing membrane vesicles excreted from *Vibrio alginolyticus* mutants lacking a polar-flagellar filament. J. Biochem. 123, 1169–1173

