## Supplemental Figure and Table for "Functional analysis of the N-terminal region of *Vibrio* FlhG, a MinD-type ATPase in flagellar number control"

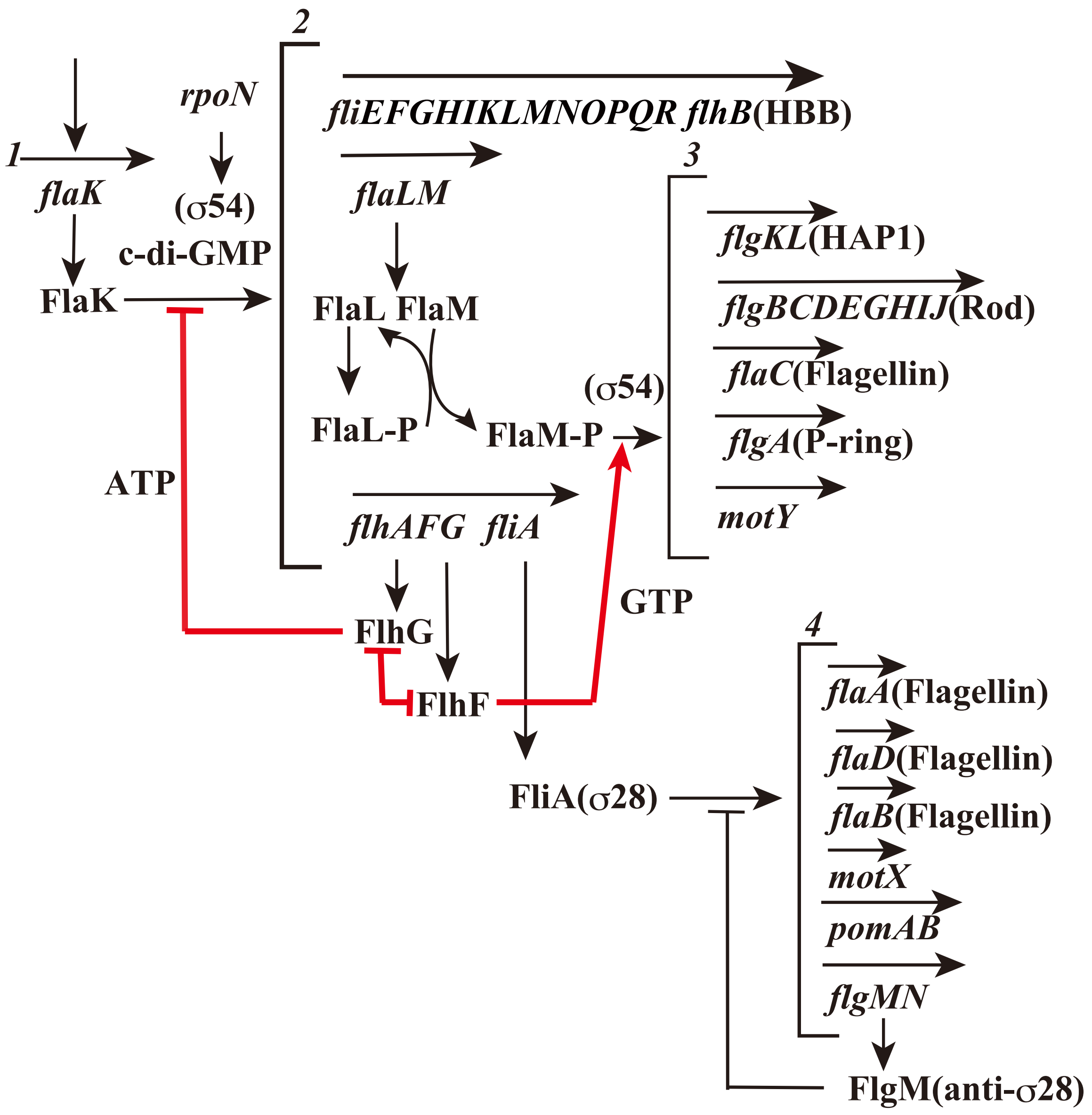

**Fig. S1.** Diagram showing the class of transcriptional control of flagellum-related factors in *V. alginolyticus*. FlhF and FlhG belong to class 2, FlhF promotes expression of class 3 gene, and FlhG inhibits the expression of class 2 genes by FlaK, which is a master regulator of flagellum-related factors.

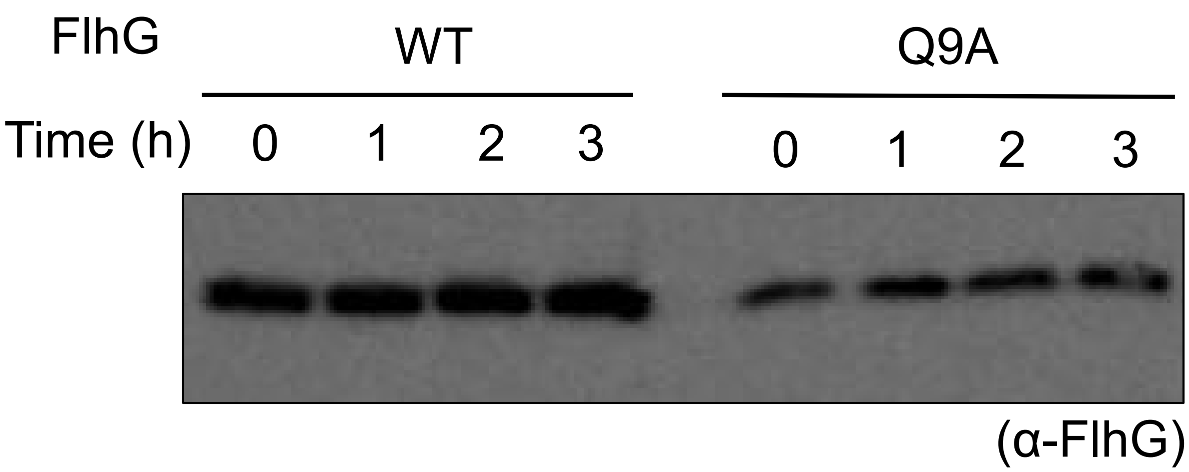

**Fig. S2.** Protein stability of the FlhG mutants. KK148 containing pAK520 encoding wild-type *flhG* (WT) or Q9A mutant (Q9A) was cultured at 30 °C in a VPG medium containing 0.02% arabinose. After 4 h, 1 mg/mL of kanamycin was added to the culture medium, and samples of the cells were collected every hour. The proteins in the cells were separated by SDS-PAGE, and the blotted membranes were probed with an anti-FlhG antibody.

**Table S1.** Bacterial strains and plasmids used in this study

| Strain or plasmid | Genotype or description | Reference or source |
|---|---|---|
| V. alginolyticus | | |
| VIO5 | VIK4 (Rif$^r$ Pof$^+$ Laf$^-$) | (1) |
| KK148 | VIO5 *flhG* | (2) |
| NMB343 | VIO5 *flhF-egfp* | This study |
| NMB363 | VIO5 Δ*flhG* (Rif$^r$ Pof$^+$ Laf$^-$ multi Pof) | This study |
| LPN1 | VIO5 Δ*flhF* (Rif$^r$ Pof$^+$ Laf$^-$) | (3) |
| LPN2 | VIO5 Δ*flhFG* (Rif$^r$ Pof$^+$ Laf$^-$) | (3) |
| E. coli | | |
| DH5α | Recipient for DNA manipulation | |
| BL21(DE3) | expression host | Novagen |
| Plasmids | | |
| pBAD33 | Cm$^r$, $P_{BAD}$ | (4) |
| pAK322 | *flhF (wt)* in pBAD33 | (2) |
| pAK325 | *flhF-egfp (wt)* in pBAD33 | (3) |
| pTSK110 | pColdIV-*flhF-his6* | (5) |
| pAK520 | *flhG* in pBAD33 | (2) |
| pAK541 | *flhG-egfp* in pBAD33 | (3) |
| pTrc-*flhG* | *$his_6$-tev-flhG* in pTrcHisB | (6) |
| pTSK29 | Cm$^r$, $P_{BAD}$, MCS of oBAD24 | (7) |
| pTSK151 | *flhG* (*wt*) in pTSK29 | This study |

Cm$^r$, chloramphenicol-resistant; Rif$^r$, Rifampicin-resistant; Pof$^+$, possessing a polar flagellum; Laf$^-$, lack of lateral flagella

(1) Okunishi et al., (1996) *J. Bacteriol.* **178**, 2409-2415. (2) Kusumoto et al., (2006) *J Biochem (Tokyo)* **139**, 113-121. (3) Kusumoto et al., (2008) *Microbiology* **154**, 1390-1399. (4) Guzman et al., (1995) *J. Bacteriol.* **177**, 4121-4130. (5) Kondo et al., (2018) *Scientific reports* **8**, 12115. (6) Ono et al., (2015) *Mol. Microbiol.* **98**, 130-141. (7) Inaba et al., (2017) *Genes Cells* **7**, 619-627.
